# Auxin confers protection against ER stress in *Caenorhabditis elegans*

**DOI:** 10.1101/2020.11.15.383760

**Authors:** A. Bhoi, F. Palladino, P. Fabrizio

## Abstract

Auxins are plant growth regulators that influence most aspects of plant development through complex mechanisms. The development of an auxin-inducible degradation (AID) system has enabled rapid, conditional protein depletion in yeast and cultured cells. More recently, the system was successfully adapted to *C. elegans* to achieve auxin-dependent degradation of targets in all tissues and developmental stages. Whether auxin treatment alone has an impact on nematode physiology is an open question. Here we show that indole-3-acetic acid (IAA), the auxin most commonly used to trigger AID in worms, functions through the conserved IRE-1/XBP-1 branch of the Unfolded Protein Response (UPR) to promote resistance to Endoplasmic Reticulum (ER) stress. Because of the central function played by the UPR in protein folding, lipid biosynthesis and lifespan regulation, these results suggest that extreme caution should be exercised when using the AID system to study these and related processes.

## Introduction

Auxins are a family of plant hormones that control gene expression during many aspects of growth and development (Teale et al., 2006). Indole-3-acetic acid (IAA), the most common natural auxin in plants, is produced mainly from tryptophan, an essential amino acid for all animals. Importantly, free-living bacteria, as well as bacterial flora in the gut of animals, can also degrade tryptophan to yield indole or indole-based compounds (Bansal et al., 2010; Lee and Lee, 2010; Lee et al., 2015).

Indoles from commensal microbiota have been shown to extend healthspan of diverse organisms, including *Caenorhabditis elegans*, *Drosophila melanogaster*, and mice, with only a negligible effect on maximal lifespan (Sonowal et al., 2017). In worms and flies, the effects of indoles on healthspan depend upon the aryl hydrocarbon receptor (AhR; also known as dioxin receptor). In *C. elegans*, transcriptional analysis showed that indole and its derivatives are important signalling molecules during bacteria-nematode interactions (Lee et al., 2017).

Whether treatment of *C. elegans* with exogenous indoles has an impact on animal physiology has not been explored. This has important implications when using the recently developed auxin-inducible degradation system (AID) that depends on the ability of IAA and other auxins to bind to the F-box transport inhibitor response 1 (TIR1) protein. TIR1 is the *Arabidopsis*-specific substrate-recognition component of the conserved SKP1-CUL1-F-box (SCF) E3 ubiquitin ligase complex, and carries out its function only in the presence of auxin. Once bound to auxin, exogenous TIR1 targets AID-tagged proteins for ubiquitin-dependent proteasomal degradation, allowing for highly efficient, conditional protein depletion in many systems (Camlin and Evans, 2019; Daniel et al., 2018; Holland et al., 2012; Lee and Lee, 2010; Natsume et al., 2016; Nishimura et al., 2009; Trost et al., 2016), including *C. elegans* (Martinez et al., 2020; Zhang et al., 2015).

In the course of our studies to investigate the tissue specific requirement for chromatin associated Heterochromatin Protein 1 in promoting resistance to ER stress in *C. elegans* (Kozlowski et al., 2014), we discovered that exposure of animals to auxin significantly increases resistance to stress of the Endoplasmic Reticulum (ER), both throughout development and in adults. The ER constitutes the entry into the secretory pathway and contributes to the maintenance of cellular calcium homeostasis, lipid synthesis, and transmembrane protein folding, making the maintenance of ER homeostasis an important component of animal physiology (Hetz, 2012). Disruptions of ER homeostasis through the accumulation of misfolded proteins in the ER by physiological, chemical, and pathological factors, also known as ER stress, activates the Unfolded Protein Response (UPR), whose role is to re-establish ER homeostasis and promote survival (Metcalf et al., 2020). We found that increased resistance to ER stress in the presence of auxin is dependent on XBP-1 and IRE-1, upstream components of the UPR, showing that auxin acts through this stress pathway to alter animal physiology. Because of the tight link between the ER stress response and essential cellular processes such as protein folding, ageing and lipid metabolism, our data suggests that auxin treatment alone may influence the outcome of experiments aimed at studying these or related processes in *C. elegans*, and possibly in other species.^1^

## Materials and Methods

### Nematode maintenance and strains

Animals were maintained under standard culture conditions (Brenner, 1974). The wildtype N2 (Bristol) was used as the reference strain. Strains used were obtained from the CGC and are listed below:

ZD417 *xbp-1(tm2457)* III; SJ17 *xbp-1(zc12)* III; RB925 *ire-1(ok799)* II; RB772 *atf-6(ok551)* X; RB545 *pek-1(ok275)* X.

### Developmental ER stress assay

4-6 day-1 adult worms were allowed to lay eggs on nematode growth medium plates (NGM) containing either tunicamycin only (Enzo Life-Sciences; 2-3 *μ*g/mL) or tunicamycin and auxin (IAA, indole-3-acetic acid, Sigma-Aldrich; 1 mM), and seeded with OP50-1 bacteria. After 3-4 hours adult worms were removed and the total number of eggs laid on each plate was counted. L4-adult worms were scored after 4 days at 20°C. 6 plates were used for each condition. Tunicamycin and IAA stock solutions were prepared in pure DMSO and ethanol (100%), respectively. DMSO and ethanol were used in control plates at the same percentage used in the experimental conditions (up to 0.03 % for tunicamycin and 0.25 % for ethanol).

Statistics and significance calculations were performed by unpaired Student’s t-test, one-way ANOVA with Tukey’s multiple comparison test, or multiple t-test with Holm-Sidak correction, as detailed in legends.

### Adult ER stress assay

The adult ER stress assay was performed as described previously (Taylor and Dillin, 2013). Briefly, worms were synchronized by bleaching and allowed to reach adulthood on NGM plates seeded with OP50-1. Day-1 adults were transferred on plates containing tunicamycin (40 *μ*g/ml) with or without auxin (1 mM). Survival was scored every day until all the animals were dead. Animals that crawled off the plates were not included in the analysis. P values were calculated using the log-rank (Mantel-Cox) method. Statistics and significance calculations for individual lifespan studies were determined using the Oasis online software (Yang et al., 2011).

## Results and Discussion

Tunicamycin (TM) induces the ER stress response by inhibiting glycosylation, leading to the accumulation of unglycosylated proteins in the ER (Travers et al., 2000). In worms, treatment with TM results in a severe developmental delay and lethality, with animals arresting at various stages of larval development (Shen et al., 2001; Shen et al., 2005). In order to assess the applicability of the AID system for studies of the ER stress response, we carried out control experiments in which we scored the impact of auxin (IAA) on survival of wildtype animals exposed to TM. Adult animals were transferred on plates containing 3 *μ*g/mL TM in the presence or absence of 1 mM auxin and the ability of progeny to reach L4-adulthood was scored after 4 days. Addition of auxin significantly improved survival, with 50-60% of animals developing to adulthood compared to 10-30% on TM alone (Figure 1, 3B). Dose response experiments with auxin concentrations ranging from 0.1 to 1 mM in the presence of 3 *μ*g/mL TM revealed increased survival with increasing auxin concentrations (Figure 1B). To test whether auxin exerts its effect uniquely throughout development, or also in adults, we performed survival assays on day 1-adult worms transferred on plates containing 40 *μ*g/mL TM in the presence or absence of 1 mM auxin. A modest but significant 10% mean survival extension was observed in the presence of auxin, indicating that the protective effect of auxin against ER stress is not limited to developmental stages (Figure 2 and S1).

**Figure 1.**
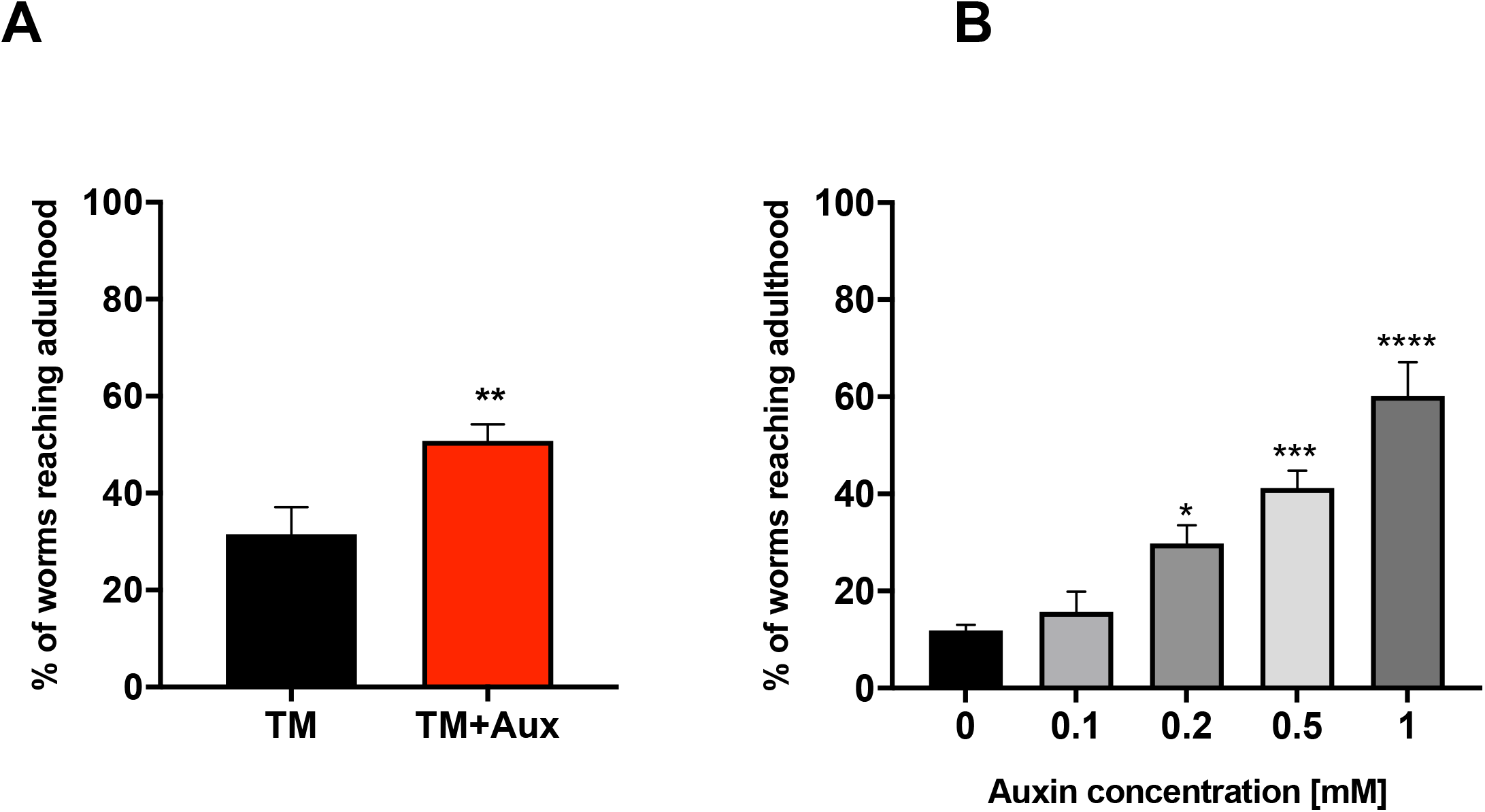
Auxin protects developing worms against ER stress. (A) Proportion of wildtype animals reaching the L4-adult stage after 4 days of development on plates containing tunicamycin (3 *μ*g/mL) with or without auxin (1 mM). Error bars represent SEM from three independent experiments (Student’s t-test, \*\**P*<0.01 auxin *vs* no auxin). (B) Proportion of wildtype animals reaching the L4-adult stage after 4 days on plates containing tunicamycin (3 *μ*g/mL) combined with increasing concentrations of auxin (0.1-1 mM). Error bars show SEM from three independent experiments (one-way ANOVA with Tukey’s multiple comparison test, \**P*<0.05, \*\*\**P*=0.0005, \*\*\*\**P*<0.0001 auxin *vs* no auxin).

**Figure 2.**
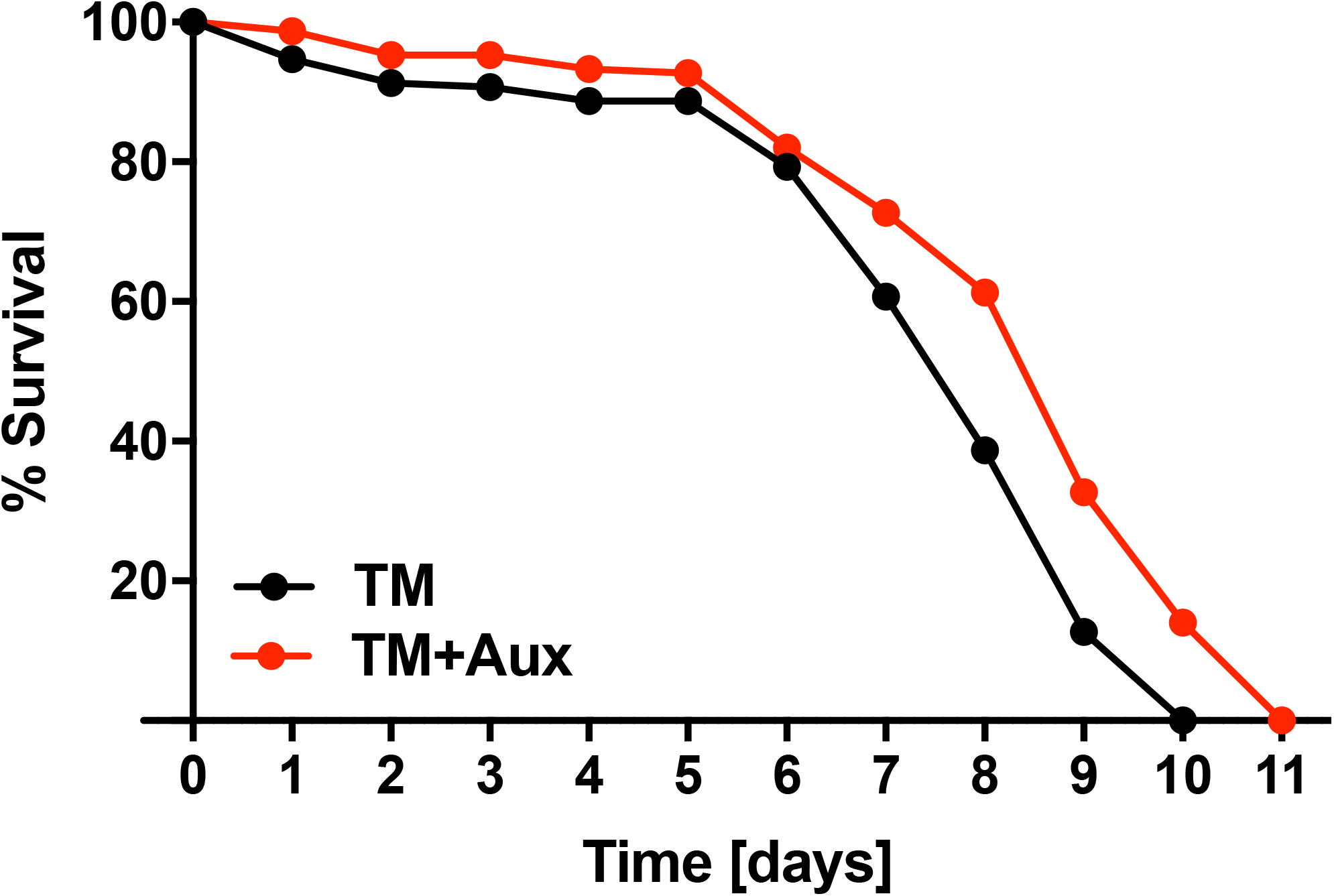
Auxin promotes ER stress resistance in adult worms. Survival of adult wildtype animals on plates containing either tunicamycin (40 *μ*g/mL) alone (N=147), or tunicamycin and auxin (1 mM) (*P*<0.0001) (N=144). Worms were exposed to auxin and/or tunicamycin starting from the first day of adulthood (day 0). The P value was calculated using the log-rank (Mantel-Cox) method. Mean and maximum lifespan were 7.8 and 10, and 8.9 and 11, for tunicamycin and tunicamycin/ auxin, respectively. A replicate experiment is shown in figure S1.

**Figure 3.**
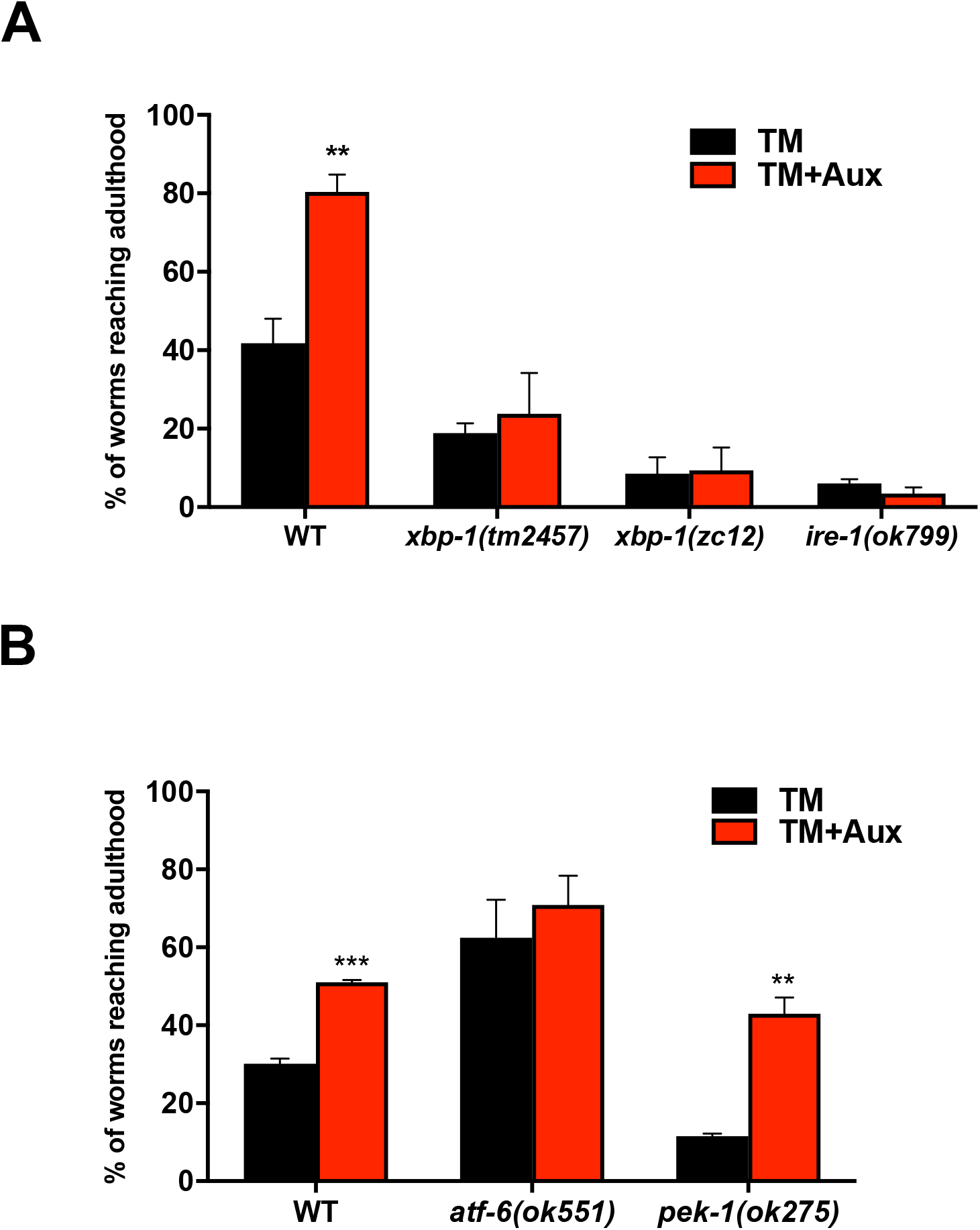
The XBP-1/IRE-1 pathway of the Unfolded Protein Response is required for auxin to induce ER stress resistance. (A) Proportion of *xbp-1(tm2457)*, *xbp-1(zc12)*, and *ire-1(ok799)* mutants, and wildtype animals reaching the L4-adult stage on plates containing either tunicamycin (2 *μ*g/mL) only or both tunicamycin and auxin (1 mM). (B) Proportion of *atf-6(ok551)* and *pek-1(ok275)* mutants, and wildtype worms reaching the L4-adult stage in presence of tunicamycin (3 *μ*g/mL) and auxin (1 mM) or tunicamycin only. For both (A) and (B) error bars show SEM from three independent experiments and statistical analysis was performed by multiple t-test with Holm-Sidak correction, \*\**P*<0.005, \*\*\**P*<0.0005 auxin *vs* no auxin).

Disruption of ER function due to ER stress activates the Unfolded Protein Response (UPR), whose role is to re-establishes ER homeostasis and promote survival via the upregulation of ER chaperones and components of ER-associated degradation (ERAD), ER expansion, and translational attenuation. In *Caenorhabditis elegans*, as in humans, three proteins sense ER stress and activate the UPR: the ribonuclease inositol-requiring protein-1 (IRE-1), the PERK kinase homolog PEK-1, and activating transcription factor-6 (ATF-6). Upon activation of the UPR in *C. elegans* the IRE-1/XBP-1 branch directs the majority of transcriptional regulation in response to acute ER stress, with PEK-1 and ATF-6 playing only minor role (Shen et al., 2005).

To test whether auxin protects animals from ER stress through the IRE-1/XBP-1 pathway, we carried out ER stress survival assays on *ire-1(ok799)*, *xbp-(tm2457),* and *xbp-1(zc12)* mutant animals on TM plates with or without auxin (Figure 3A). *ire-1(ok799)* is a null mutation (Roux et al., 2016), *xbp-1(tm2457)* is a deletion at the 3’ end of the gene (Levi-Ferber et al., 2015), and *xbp-1(zc12)* a stop codon at the 5’ end of the gene (Calfon et al., 2002). Mutants carrying these alleles are extremely sensitive to TM (figure 3A), as previously shown (Calfon et al., 2002; Henis-Korenblit et al., 2010; Roux et al., 2016); a lower concentration of TM, 2 *μ*g/mL, was therefore used to allow the detection of at least 10-20 L4/adults on the TM plates. Addition of auxin improved neither developmental defects, nor survival in the presence of TM (figure 3A). The stronger effect of TM on *xbp-1(zc12)* compared to *xbp-1(tm2457)* suggests that *xbp-1(tm2457)* may be a strong hypomorph, rather than a null allele. Therefore, auxin acts through the IRE-1/XBP-1 pathway to confer enhanced ER stress resistance. By contrast, further analysis showed that the putative null allele *pek-1(ok275)* (Henis-Korenblit et al., 2010) did not abolish the protective effect of auxin: these animals were as resistant to ER stress as wildtype, ruling out a major role for the PEK-1 branch of the UPR in mediating the effect of auxin on ER stress resistance. Interestingly, *atf-6(ok551)* putative null mutants (Henis-Korenblit et al., 2010) showed a strong increase in resistance to TM compared to wildtype worms, and auxin treatment only resulted in a marginal further increase (figure 3B). Protection against ER stress in the absence of *atf-6* has been previously described, but the mechanisms behind this effect remain elusive (Bischof et al., 2008; Burkewitz et al., 2020; Shen et al., 2005)

Together, our results point to the IRE-1/XBP-1 branch of the UPR as the main mediator of the effect of auxin on the ER stress response. These results also rule out the possibility that auxin acts by preventing the uptake of ER stress inducing drugs such as TM.

Overall, the data included in this communication show that exposure of *C. elegans* to auxin (IAA) protects animals from ER stress, both during development and as adults. Increased resistance to ER stress by auxin is dependent on the canonical UPR, including upstream components IRE-1/XBP-1. These results suggest that exogenous indoles can impact *C. elegans* physiology.

Indole closely resembles human and plant hormones such as serotonin and IAA, leading to the speculation that it is the archetype for cell hormones (Tomberlin et al., 2017). Our findings are consistent with recent data showing that indole and its derivatives are important signalling molecules during bacteria-nematode interactions. *C. elegans* can sense and moves towards indole and indole-producing bacteria, but avoids non-indole producing pathogenic bacteria (Lee et al., 2017). Furthermore, indole-producing and non-indole-producing bacteria exert divergent effects on *C. elegans* egg-laying behaviour, and various indole derivatives also increase chemotaxis and egg-laying al low concentrations (Lee et al., 2017).

The effect of indoles and indole derivates has been also tested on nematode lifespan. Indole at a concentration range of 0.1 – 0.25 mM was found to promote healthy aging (Sonowal et al., 2017) while at concentrations higher than 0.5 mM it was found to reduce survival (Lee et al., 2017). By contrast, 1 mM IAA was reported to have no effect on either lifespan or mitochondrial functions (Dilberger et al., 2020; Kasimatis et al., 2018). The reasons for these different responses when using closely-related molecules are unclear. However, the beneficial effect of indole on healthspan was also observed in *Drosophila* and mice (Sonowal et al., 2017), suggesting conserved mechanisms.

The AID system is increasingly used and constantly being improved to achieve rapid and reversible protein degradation in different systems (Martinez et al., 2020; Sathyan et al., 2019). Our results suggest that caution should be applied when using this system to study stress related pathways in *C. elegans*, and perhaps in other species. More generally, our study illustrates how indoles and related metabolites can influence various physiological processes, with potential implications for healthy and diseased states in man (Roager and Licht, 2018).

## Acknowledgements

We thank members of the Palladino Lab for helpful discussions.

---

Figure S1. Auxin induces ER stress resistance in adult worms. Repetition of the experiment shown in figure 3. Survival of adult wildtype worms on plates containing tunicamycin (40 *μ*g/mL) alone (N=153) or tunicamycin and auxin (1 mM) (*P*<0.0001) (N=154). The P value was calculated using the log-rank (Mantel-Cox) method. Mean and maximum lifespan were 8.1 and 10, and 8.6 and 11, for tunicamycin and tunicamycin/auxin, respectively.

